# Quantitative study of unsaturated transport of glycerol through aquaglyceroporin that has high affinity for glycerol

**DOI:** 10.1101/2020.06.15.152512

**Authors:** Roberto A. Rodriguez, Ruth Chan, Huiyun Liang, Liao Y. Chen

## Abstract

**Graphical Abstract:** 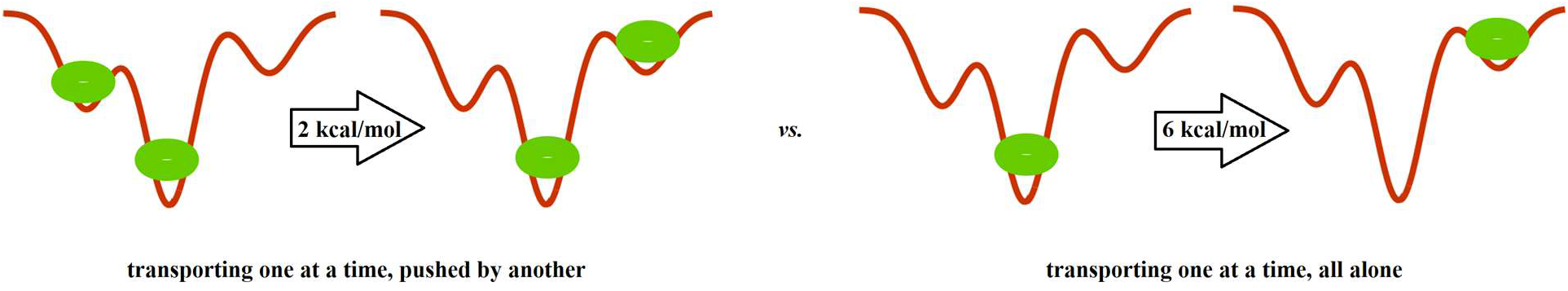

**ABSTRACT:** The structures of several aquaglyceroporins have been resolved to atomic resolution showing two or more glycerols bound inside a channel and confirming a glycerol-facilitator’s affinity for its substrate glycerol. However, the kinetics data of glycerol transport experiments all point to unsaturated transport that is characteristic of low substrate affinity in terms of the Michaelis-Menten kinetics. In this article, we present an *in silico-in vitro* research focused on AQP3, one of the human aquaglyceroporins that is natively expressed in the abundantly available erythrocytes. We conducted 2.1 μs *in silico* simulations of AQP3 embedded in a model erythrocyte membrane with intracellular-extracellular asymmetries in leaflet lipid compositions and compartment salt ions. From the equilibrium molecular dynamics (MD), we elucidated the mechanism of glycerol transport at high substrate concentrations. From the steered MD simulations, we computed the Gibbs free-energy profile throughout the AQP3 channel. From the free-energy profile, we quantified the kinetics of glycerol transport that is unsaturated due to glycerol-glycerol interaction mediated by AQP3 resulting in the concerted movement of two glycerol molecules for the transport of one glycerol molecule across the cell membrane. We conducted *in vitro* experiments on glycerol uptake into human erythrocytes for a wide range of substrate concentrations and various temperatures. The experimental data quantitatively validated our theoretical-computational conclusions on the unsaturated glycerol transport through AQP3 that has high affinity for glycerol.

## INTRODUCTION

The transport of glycerol across the cell membrane is fundamental in human physiology, biology, and biotechnology, which has been investigated since long ago [*e.g.* ^1–7^]. Aquaglyceroporins, a subfamily of membrane proteins belonging to the aquaporin (AQP) family (see, e.g., ^8–75^ for the background literature and some recent research), facilitate glycerol diffusion across the cell membrane down the concentration gradient. In the human body, there are four aquaglyceroporins: AQP3^23–27^, AQP7^62^, AQP9^34, 47, 63, 64^, and AQP10^46, 65–69^ which are responsible for lipid homeostasis and other physiological functions. In other organisms, there are, *e.g., E. coli* aquaglyceroporin GlpF^7, 20, 44, 73^ and *P. falciparum* aquaporin PfAQP^22, 45, 70^. Pathologically, for example, AQP3 found in various epithelial cells is implicated in cancers; AQP3 and AQP10 expressed in enterocytes and adipocytes are relevant in obesity research. Today, four aquaglyceroporin structures have been resolved to the atomic resolution: GlpF in 2000^20^, PfAQP in 2008^22^, AQP10 in 2018^69^, and AQP7 in 2020^60, 76^. All of these high-resolution X-ray structures showed glycerols bound inside the channels, demonstrating that aquaglyceroporins have affinity for their substrate, glycerol. Kinetics experiments on AQP9 and AQP10 showed saturated transport at low substrate concentrations^46, 47^. However, many other kinetics experiments of glycerol transport showed unsaturable characteristics for glycerol concentrations up to ~1 M^5, 6, 71–73^, which, based on the Michaelis-Menten formalism, would suggest that aquaglyceroporins are simple channels without affinity for their substrate. Putting all these together, we have a paradox: Are aquaglyceroporins facilitators with high affinity for their substrate or simple channels propping a pore for glycerol passage through the cell membrane without much affinity for the substrate?

In this paper, we present an *in silico-in vitro* investigation of AQP3 in the erythrocyte membrane aimed at resolving this outstanding paradox: How can an aquaglyceroporin have high affinity for glycerol but also transport the substrate in an unsaturable manner? Our study is based on the following experimentally validated characteristics: Each human erythrocyte carries approximately 2,000 copies^77^ of AQP3 that are responsible for glycerol transport across the cell membrane^25, 28^. The membrane lipid compositions of human erythrocytes^78–82^ are approximately 11% phosphatidylcholine (POPC), 38% phosphatidylethanolamine (POPE), 22% phosphatidylserine (POPS), 9% sphingomyelin (SSM), and 20% cholesterol (CHL) in the inner leaflet and 35% POPC, 10 POPE, 35% SSM, and 20% CHL in the outer leaflet, respectively.

We first ran equilibrium molecular dynamics (MD) simulations for 16.0 monomer μs (0.5 μs for a large system having 32 AQP3 monomers, shown in Fig. 1) which is sufficiently long for direct observation of a few glycerol transport events through AQP3. In an event of transporting a substrate across the cell membrane, a glycerol molecule can bind to AQP3 from one side of the membrane and dissociate from AQP3 to the other side. This pathway dominates when the substrate concentration is low and the probability is negligible for simultaneous binding of multiple substrates. Our large-scale simulation (having 200 mM glycerol in the system) showed a new transport pathway. Along this pathway, two substrates can bind simultaneously inside the AQP3 channel, lining up in a single file manner. The two substrates bound inside the channel next to each other are in constant thermal fluctuations pushing one another and thus making it easy for one of them to dissociate from the protein to complete the transport of one glycerol molecule. This new pathway can be exemplified in a glycerol efflux event: There is always a glycerol bound at the high affinity site near the center of the channel. A second glycerol binds to a low affinity site on the intracellular (IC) side next to the first glycerol. When this second glycerol falls into the high affinity site, it pushes the first one to the low affinity site on the extracellular (EC) side. From there, the first glycerol can easily dissociate from AQP3 into the EC fluid. This second transport pathway would become more significant as the substrate concentration increases, resulting in unsaturable transport characteristics of an aquaglyceroporin. It is interesting to note that, long before the discovery of aquaporins, it was already proposed^3, 6^ that glycerol transport involved simultaneous binding of two glycerol molecules from one side of the membrane and releasing them on the other side membrane. Now, we have the atomistic details about the inner workings of an aquaglyceroporin.

**Fig. 1.**
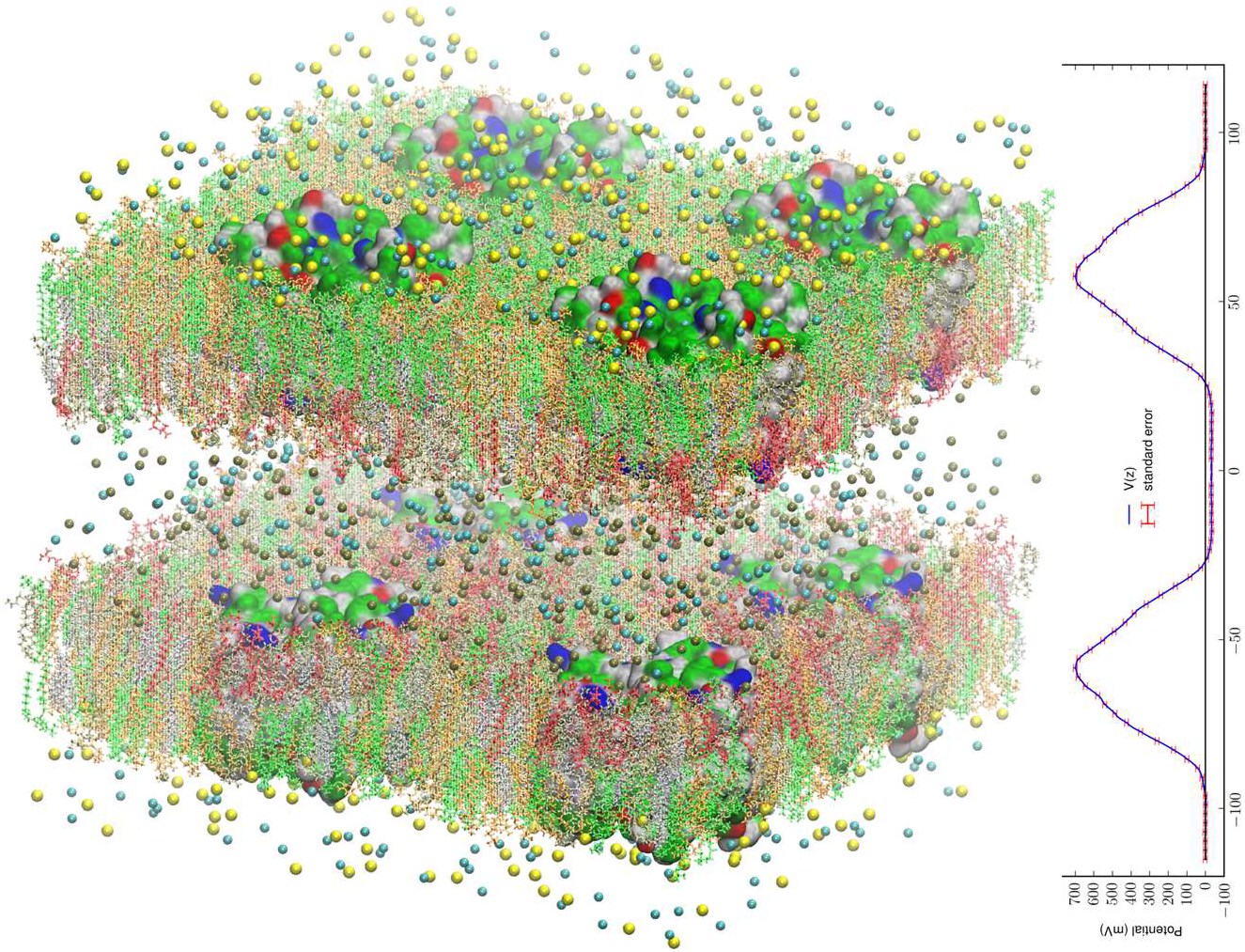
All-atom model system of eight AQP3 tetramers in asymmetric environments mimicking the membrane of a human erythrocyte. The system consists of 1,249,096 atoms. The intracellular space is located at −40Å < *z* < 40Å and the extracellular space at *z* < −95Å and *z* > 95Å. The water and glycerol molecules are not shown for clearer views of all the other constituents of the system. The intracellular salt (KCl) and the extracellular salt (NaCl) are shown as spheres colored by atom names (Cl, cyan; K, metallic; Na, yellow). The lipids are represented as licorices colored by lipid names (POPC, orange; POPE, tan; POPS, red; SSM, green; CHL, silver). The proteins are presented as surfaces colored by residue types (hydrophilic, green; hydrophobic, white; positively charged, blue; negatively charged, red). Molecular graphics in this paper were rendered with VMD^83^. The PME-based electrostatic potential is shown in the right panel.

For quantitative characteristics, we ran 1.68 μs steered MD to map out the driving force, namely the gradient of the potential of mean force (PMF) that is the Gibbs free energy of the system as a function of the coordinates of a glycerol molecule along the transport path throughout AQP3 between the cytoplasm and the EC fluid. From the PMF curve and the fluctuations at three binding sites inside the AQP3 channel, we computed the glycerol-AQP3 affinities showing high affinity ~500/M at the central binding site near the Asparagine-Proline-Alanine (NPA) motifs and low affinities ~1/M at two other sites, one on the IC side and one on the EC side. We further quantified the transport kinetics based on the PMF curve to predict the time courses of erythrocyte swelling-shrinking caused by glycerol uptake. We conducted *in vitro* experiments of glycerol uptake for five substrate concentrations ranging from 25 mM to 400 mM at four temperatures from 5°C to 37ºC. The experimental data invariably validated the computational predictions with a single fitting parameter that is the rate constant for glycerol to bind on to AQP3 from the IC side.

## METHODS

The parameters, the coordinates, and scripts for setting up the model systems, running the simulations, and analyzing the data are available at Harvard Dataverse via https://doi.org/10.7910/DVN/V9I4YQ.

### Model system setup and simulation parameters

Following the well-tested steps of the literature, we employed CHARMM-GUI^84^ to build an all-atom model of an AQP3 tetramer embedded in a 117Å×117Å patch of erythrocyte membrane. The AQP3 coordinates were taken from Ref. ^53^ (optimized homology model validated with *in vitro* experiments). The positioning of the AQP3 tetramer was determined by matching the hydrophobic side surface with the lipid tails and aligning the channel axes perpendicular to the membrane. The AQP-membrane complex was sandwiched by two layers of TIP3P waters, each of which was approximately 30Å thick. The system was then neutralized and salinated with Na^+^ and Cl^−^ ions to a salt concentration of 150 mM. Glycerol was added to the system to 200 mM in concentration. The system so constructed consists of four AQP3 monomers in a single patch of membrane constituted with 156,137 atoms, which is referred to as SysI (shown in Supplemental Information, SI, Fig. S1). We employed NAMD 2.13^85^ as the MD engine. We used CHARMM36 parameters^86–88^ for inter- and intra-molecular interactions. After the initial equilibration steps, we fully equilibrated the system by running unbiased MD for 500 ns with constant pressure at 1.0 bar (Nose-Hoover barostat) and constant temperature at 303.15 K (Langevin thermostat). The Langevin damping coefficient was chosen to be 1/ps. The periodic boundary conditions were applied to all three dimensions. The particle mesh Ewald (PME) was used for the long-range electrostatic interactions (grid level: 128×128×128). The time step was 2.0 fs. The cut-off for long-range interactions was set to 10 Å with a switching distance of 9 Å.

We replicated the fully equilibrated SysI seven times to obtain eight copies of SysI. With appropriate translations and rotations of the eight copies, we formed SysII, a large system of two patches of membrane that consists of 1,249,096 atoms. In each patch of the membrane, there are four AQP3 tetramers (16 monomers). We replaced Na^+^ ions in the intracellular space with K^+^ ions. In this manner, we constructed an all-atom AQP3-membrane system (illustrated in Fig. 1) that has an intracellular saline of KCl separated by two membrane patches from the extracellular saline of NaCl. The model membrane has the asymmetry in the lipid compositions of the inner and the outer leaflets mimicking the human erythrocyte membrane. Unbiased MD was run for 500 ns for this large SysII with identical parameters used for SysI except that the PME was implemented on a grid level of 256×256×256. The last 200 ns of the trajectories were used in the computation of the membrane potential (Fig. 1) and the search for the glycerol transport pathways. The advantages of SysII are twofold. First, the 200 ns production run of this large system having 8 copies of the AQP3 tetramer is statistically equivalent to a 1.6 μs run of SysI that has only one biologically functional unit (AQP3 tetramer). This statistical time scale is more than the average time, 1.1 μs, for a single transport event per AQP3 tetramer. A sub-microsecond run of SysII, which is achievable in days, not weeks, on high-performance supercomputers, should be sufficient for the search of transport mechanisms. Second, SysII has a significant enhancement of the signal to noise ratio over SysI because the intrinsic pressure fluctuation of a system is inversely proportional to the volume of the system^89^. This second advantage, while not critical in this study, may be needed in other studies where the biophysical processes are sensitive to the pressure fluctuations.

### Computing the Gibbs free-energy profile

We conducted 1,680 ns steered MD of SysI (illustrated in Fig. 2 and SI, Fig. S1) to compute the PMF along the glycerol transport path through an AQP3 channel across the membrane. We followed the multi-sectional protocol detailed in Ref. ^75^. We defined the forward direction as along the z-axis, from the EC side to the IC side. We divided the entire glycerol transport path across the membrane from *z* = −20Å to *z* = 20Å into 40 evenly divided sections. From the central binding site (*z* = 0Å, shown in Fig. 2) to the IC side (*z* ≥ 20Å), the center-of-mass z-degree of freedom of glycerol was steered at a speed of 0.25 Å/ns for 4 ns over one section for a z-displacement of 1.0 Å to sample a forward path over that section. At the end of each section, the z-coordinate of the glycerol center-of-mass was fixed (or, technically, pulled at a speed of 0.0 Å/ns) while the system was equilibrated for 10 ns. From the end of the 10 ns equilibration, the z-coordinate of the glycerol center-of-mass was pulled for 4 ns for a z-displacement of −1.0 Å to sample a reverse path. From the binding site (*z* = 0Å) to the EC side (*z* ≤ −20Å), the center-of-mass z-degree of freedom of glycerol was steered for 4 ns for a z-displacement of −1.0 Å to sample a reverse path over one section. At the end of that section, the z-coordinate of the glycerol center-of-mass was fixed while the system was equilibrated for 10 ns. From the end of the 10 ns equilibration, the z-coordinate of the glycerol center-of-mass was pulled for 4 ns for a z-displacement of +1.0 Å to sample a forward path. In this way, section by section, we sampled a set of four forward paths and four reverse paths in each of the 40 sections (20 sections from the central binding site to the IC side and 20 sections from the central binding site to the EC side) along the entire transport path between the EC and the IC sides. The force acting on the glycerol center-of-mass was recorded along the forward and the reverse pulling paths for computing the PMF along the entire transport path from the EC side to the central binding site and then to the IC side. The PMF was computed from the work along the forward paths and the work along the reverse paths via the Brownian-dynamics fluctuation-dissipation theorem^75^.

**Fig. 2.**
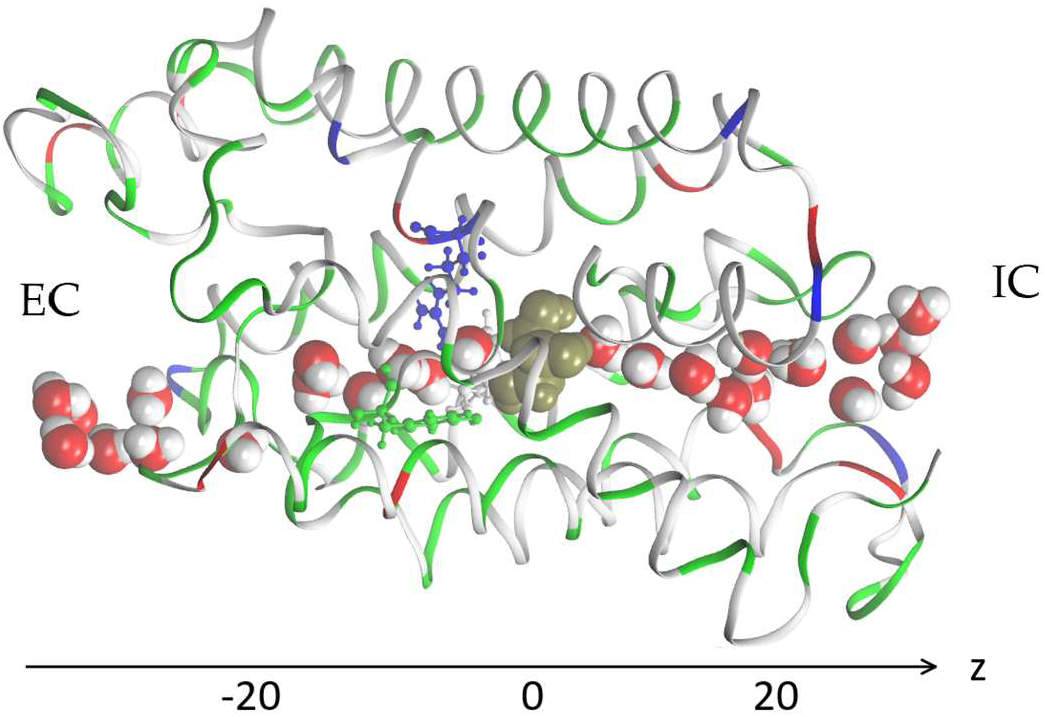
AQP3 monomer channel with a glycerol molecule (gold colored spheres) at the central binding site near the NPA motifs. The whole monomer protein is shown as ribbons along with the aromatic/Arginine motif residues (Phe63, Tyr212, and Arg218) are shown in ball-and-sticks, all colored by residue types (See Fig. 1 for colors). The water molecules inside and near the channel are shown in space-filling spheres colored by atoms (O, red; H, white).

### Binding affinity from PMF in 3D

Following the standard literature (*e.g.*, ^90^), one can relate the binding affinity (inverse of the dissociation constant k_Di_) at the *i*-th binding site to the PMF difference in 3 dimensions (3D) and the two partial partitions as follows:

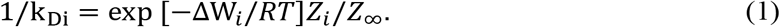

Here Δ*W*_*i*_ is the PMF at the *i*-th binding site minus the PMF in the dissociated state when glycerol is far away from the protein. *R* is the gas constant. *T* is the absolute temperature. *Z*_*i*_ is the partial partition of glycerol in the *i*-th bound state which can be computed by sampling the fluctuations in 3 degrees of freedom of the glycerol center of mass and invoking the Gaussian approximation for the fluctuations in the bound state^91^.

### Experimental assays and analyses

The packed red blood cells from three anonymous healthy donors were purchased from the South Texas Blood and Tissue Center. Equal volumes of the three erythrocyte samples were mixed and washed three times and then suspended in phosphate buffered saline (PBS) to a total concentration of 4% hematocrit. In each experimental run, 5 μL of the 4% hematocrit suspension in PBS was rapidly mixed with an equal volume of PBS containing glycerol at a concentration of 2Δ*c*^*GOL*^ using an Applied PhotoPhysics SX20 stopped-flow spectrometer. In such a mixture solution, the extracellular environment of the erythrocyte is initially hyperosmotic with a gradient of Δ*c*^*GOL*^which drives a water efflux out of the cell resulting in cellular shrinkage in the initial phase. However, the extracellular glycerol diffuses through AQP3 into the cytoplasm. Subsequent to the initial phase, the glycerol influx drags along a water influx resulting in cellular swelling which lasts until the glycerol gradient dissipates away and the cells return to the state before mixing. The intensity of light scattered at 90° was measured to monitor the erythrocyte shrinking-swelling process. The scattered light intensity is related to the varying cellular volume as follows:

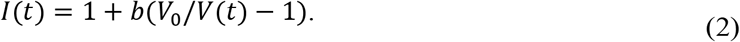

Here *t* is time. *b* is a parameter to account for the number of erythrocytes in the samples of a given set of experiments. *V*(*t*) is the intracellular volume of an erythrocyte with its initial value noted as *V*_0_. It follows the following dynamics equation:

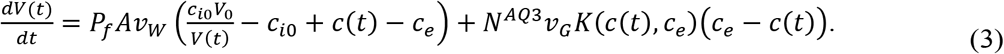

Here *P*_*f*_ is the osmotic permeability and *A* is the cellular surface area. The *P*_*f*_*A* values of the erythrocyte samples were measured in Ref. ^82^. *c*_*i*0_ is the initial concentration of impermeable solutes. *c*_*e*_ = Δ*c*^*GOL*^ is the EC concentration of glycerol, which is approximately constant because the IC spaces of all the erythrocytes together amount to no more than 2% of the EC space in the mixture solution of an experiment run. *v*_*w*_ and *v*_*G*_ are the molar volume of water and glycerol respectively. *c*(*t*), the IC glycerol concentration, is determined by the glycerol transport through AQP3,

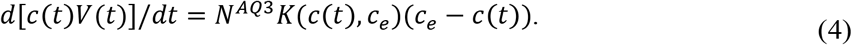

Here *N*^*AQ3*^ is the number of moles of AQP3 per erythrocyte. The flux of glycerol uptake is proportional to the glycerol concentration gradient and the coefficient *K*(*c*(*t*), *c*_*e*_) has a very complex dependence upon the IC and the EC concentrations, which is given in SI, Part I. That complex dependence reflects the dynamic balances among the glycerol molecules occupying the three binding sites inside an AQP3 channel.

For each experiment, the full dynamics in Eqs. (3) and (4) were numerically integrated for a given set of parameters (one of which was used a fitting parameter) for a time-course solution in Eq. (2), which was least-squares fitted to the measured intensity of scattered light in the cellular shrinking-swelling process.

## RESULTS AND DISCUSSION

### Mechanisms of glycerol transport

In order to obtain direct observation of a transport event, we conduct 500 ns unbiased equilibrium MD run of SysII (shown in Fig. 1) having a glycerol concentration of 200 mM. We observed that glycerol molecules bind inside an AQP3 channel at three different locations illustrated in Fig. 3. We note the central binding site near the NPA motifs as Site 0, the one on the EC side of the NPA as Site 1, and the one on the IC side of the NPA as Site 2. There are eight possible states for three binding sites being occupied or unoccupied (Table I). However, State 101 was not observed in the simulation and thus not considered in this study. Fig. 3A illustrates the seven states that were observed in the simulation. Fig. 3B illustrates three of them and the transitions between them that gave rise to the IC-to-EC transport of one glycerol molecule *via* the collective motion of two glycerol molecules.

**Table I.** States of binding occupation (G means site occupied)

| Binding site | State 000 | State 010 | State 011 | State 100 | State 110 | State 101 | State 111 | State 001 |
| --- | --- | --- | --- | --- | --- | --- | --- | --- |
| EC side, Site 1 |  |  |  | G | G | G | G |  |
| Central, Site 0 |  | G | G |  | G |  | G |  |
| IC side, Site 2 |  |  | G |  |  | G | G | G |

**Fig. 3.**
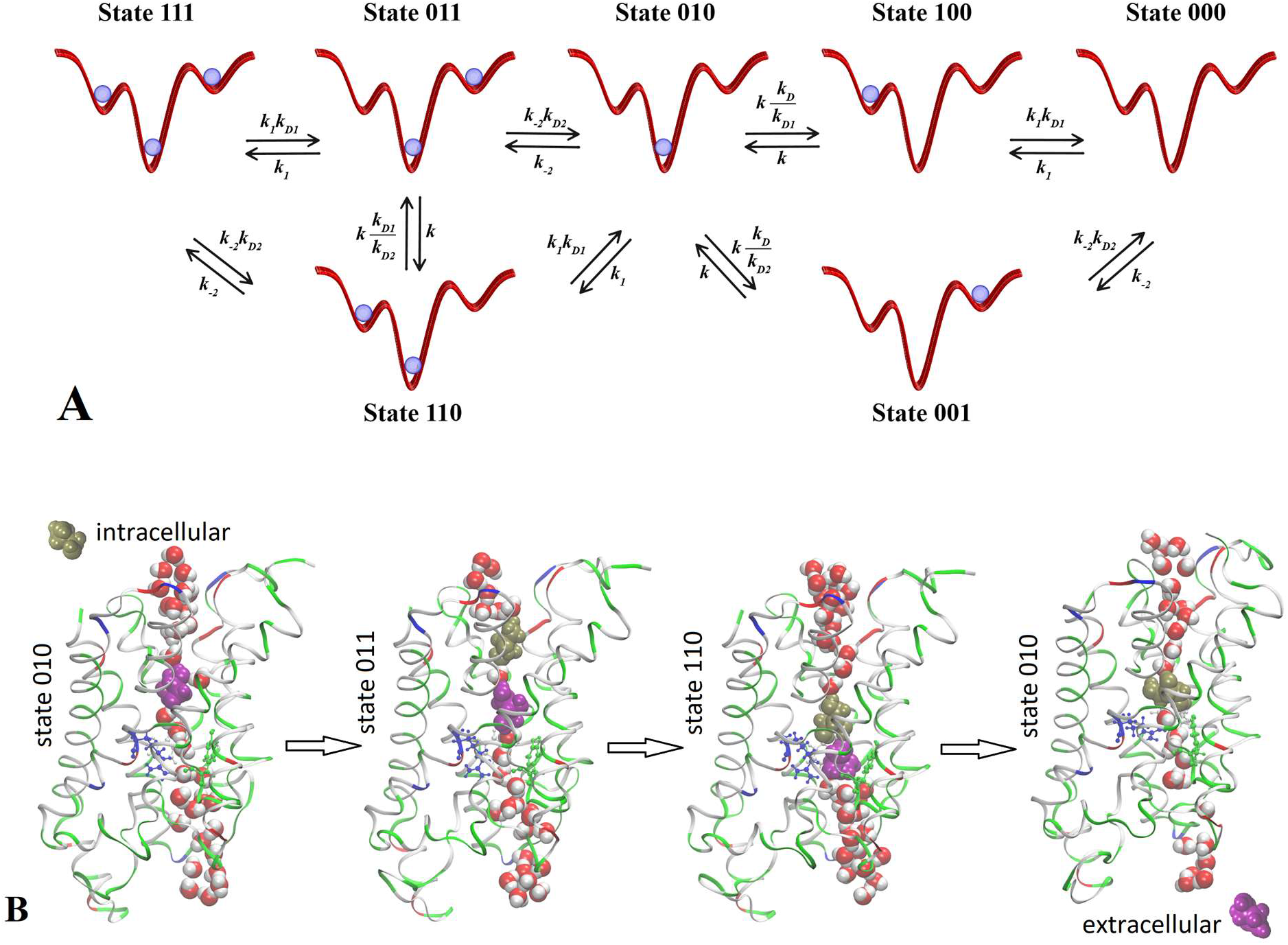
A. Seven states of the glycerol-AQP3 complex observed and rate constants of transitions between them. The three wells on a red curve indicate the binding site of glycerol inside the AQP3 channel. The blue dot indicates the presence of a glycerol molecule at the binding site. B. An IC-to-EC transport event. The protein AQP3 monomer is shown in ribbons and the ar/R sf residues (Phe63, Tyr212, Arg218) in ball-and-sticks, all colored by residue types (hydrophobic, white; hydrophilic, green; negatively charged, red; positively charged, blue). Water molecules inside and near the AQP3 channel are shown in space-filling spheres colored by atoms (O, red; H, white). Two glycerol molecules are shown in space-filling spheres: GOL661 colored purple and GOL102 colored gold.

Examining the trajectories during the last 200 ns (after SysII was fully equilibrated), we gained the following insights: Site 0 was always occupied indicating that State 010 has the lowest free energy. States 000, 100, 111, 001 were nearly never occupied indicating that either Site 1 or Site 2 has much higher free energy than Site 0. Double occupancies (State 011 and State 110) did show up, noting that SysII has a high glycerol concentration of 200 mM. The transitions between States 010 and 110 occurred multiple times and so did the transitions between States 010 and 011. These transitions indicated that the free energy for a glycerol being at Site 1 or 2 is not much lower than the bulk level (when it is outside the channel on the EC or IC side). They further indicated that there is no significant attraction between the two glycerols in State 011 or 110. Transitions between State 011 and State 110 also occurred within the 0.2 μs time frame, indicating that States 011 and 110 have similar free energies. All of these insights will be quantitatively validated when the free-energy profile is computed along the glycerol diffusion path throughout the AQP3 channel.

Qualitatively, we now are clear that there are two pathways for glycerol transport across the cell membrane through AQP3. At a low glycerol concentration, double or triple occupancy inside the channel is very low. Namely, States 110 and 011 (double occupancy) have negligibly small probabilities and even more so do States 101 (double occupancy) and 111 (triple occupancy). The EC to IC transport pathway consists of consecutive transitions 000 to 100 to 010 to 001 to 000 and the IC to EC transport pathway consists of the same transitions in reverse sequence. The bottleneck of the transport pathway is the 010 to 001 transition or 010 to 100 transition because the central binding site has high affinity. At a higher glycerol concentration, double occupancies inside the channel become more probable. A second transport pathway starts to play a more significant role. This second pathway consists of consecutive transitions 010 to 011 to 110 to 010 for a glycerol efflux event (illustrated in Fig. 3) or the same transitions in reverse order for a glycerol uptake event.

In principle, it is desirable extend the unbiased equilibrium MD to many milliseconds in length to achieve statistically significant results for glycerol transport. In practice, it is infeasible and unnecessary to incur such a great computing cost because quantitatively accurate results can be obtained from steered MD simulations for less than two microseconds.

### Quantitative characteristics

Fig. 4 shows the PMF throughout the AQP3 channel as a function of the z-coordinate of the glycerol’s center of mass. It represents the Gibbs free energy of the system when a glycerol molecule is located at a given location. The reference level of the PMF was chosen at the bulk level on either the EC or the IC side. The two bulk levels must be equal for neutral solute transport across the cell membrane which is not an actively driven process but a facilitated passive process of diffusion down the concentration gradient. The PMF curve leveling off to zero on both the EC side and the IC side in Fig. 4 indicates accuracy of our computation. Inside the protein channel, the PMF presents a deep well (Δ*W*_1_ = −7.5 kcal/mol) near the NPA motifs (around *z*~0), which is a binding site for glycerol (Site 0). On the EC side, between the NPA and the aromatic/Arginine (ar/R) selectivity filter (sf), there is another binding site (Site 1) where the PMF has a local minimum (Δ*W*_n_ = −3.5 kcal/mol). The third binding site (Site 2) is located on the IC side of the NPA where the PMF is Δ*W*_7_ = −2.2 kcal/mol. These PMF well depths of the three binding sites are the main factors to determine the affinities (the inverse dissociation constants): 1/*k*_*D*_ = *f*_0_exp [−Δ*W*_0_/*RT*] for Site 0, 1/*k*_*D*1_ = *f*_1_exp [−Δ*W*_n_/*RT*] for Site 1, and 1/*k*_*D*2_ = *f*_2_exp [−Δ*W*_7_/*RT*] for Site 2. The other factors involved to determine the affinities are the fluctuations at the three sites (*f*_0_, *f*_1_, *f*_2_) which was computed straightforwardly from the equilibrium MD runs with the gaussian approximation. Combining the fluctuations and the PMF well depths, we obtained the dissociation constants tabulated in Table II.

**Table II.** Computed dissociation constants

| | Central binding site, $k_D$ | EC binding site, $k_{D1}$ | IC binding site, $k_{D2}$ |
| --- | --- | --- | --- |
| 36.6 °C | 4.0 mM | $0.9 \times 10^3 \text{ mM}$ | $1.2 \times 10^3 \text{ mM}$ |
| 23.5 °C | 2.0 mM | $0.5 \times 10^3 \text{ mM}$ | $0.8 \times 10^3 \text{ mM}$ |
| 16.1 °C | 1.6 mM | $0.4 \times 10^3 \text{ mM}$ | $0.7 \times 10^3 \text{ mM}$ |
| 5.3 °C | 1.0 mM | $0.35 \times 10^3 \text{ mM}$ | $0.55 \times 10^3 \text{ mM}$ |

**Fig. 4.**
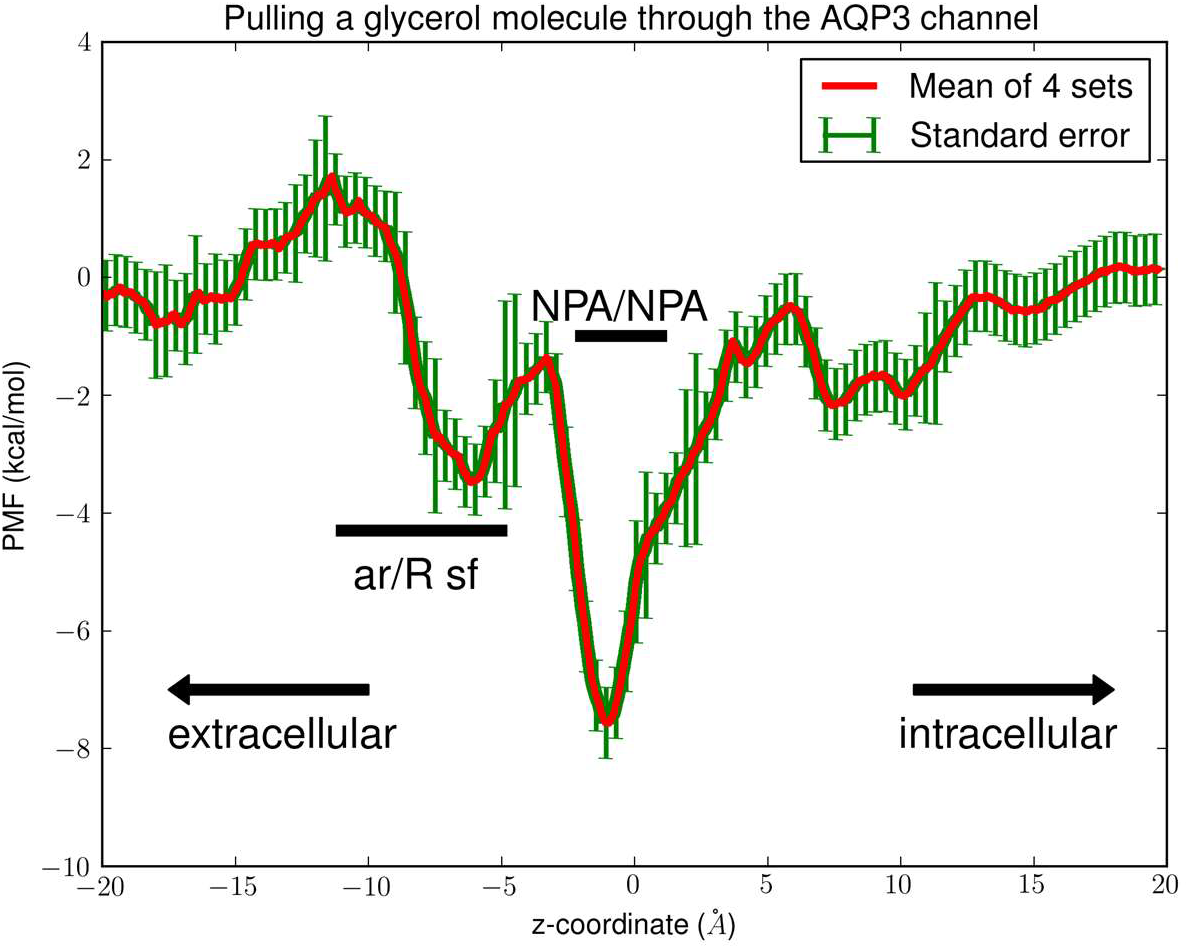
PMF of glycerol throughout the AQP3 channel. The coordinates are set so that the center of membrane is located at *z*~0Å. In the single-file region (−10Å < *z* < 10Å), the PMF is one dimensional. In the IC (*z* < −10Å) or EC (*z* > 10Å) side of the channel, the PMF is three dimensional. The three binding sites are located at: Site 1, *z*~ − 6Å; Site 0, *z*~0Å; Site 2, *z*~8Å.

Interestingly, the PMF curve dictates that the rate constants for transitions between the states are related to one another as tabulated in Table III. Additionally, *k* = *k*_−2_max (*k*_*D*1_, *k*_*D*2_) and *k*_1_ = *k*_−2_exp (−Δ*W*_*EC*_/*RT*) where Δ*W*_*EC*_ ≃ 1.8 kcal/mol is the PMF barrier along the binding path of a glycerol molecule from the EC side. *k*_n7_ is the rate constant for glycerol binding onto AQP3 from the IC side. With these relationships, we leave one rate constant *k*_−2_ free as a fitting parameter to fit with multiple sets of experimental data for five glycerol concentrations at four different temperatures. If this PMF-based theoretical-computational framework accurately captured all the fundamental steps in glycerol transport through AQP3, at each temperature, *k*_−2_ would be nearly independent of the glycerol concentration. The values of *k*_−2_ are expected to be slightly dependent on the experimental temperature because the process represented by *k*_−2_ (a glycerol molecule binding onto AQP3 from the IC side) is essentially downhill without significant barriers in the PMF profile.

**Table III.**
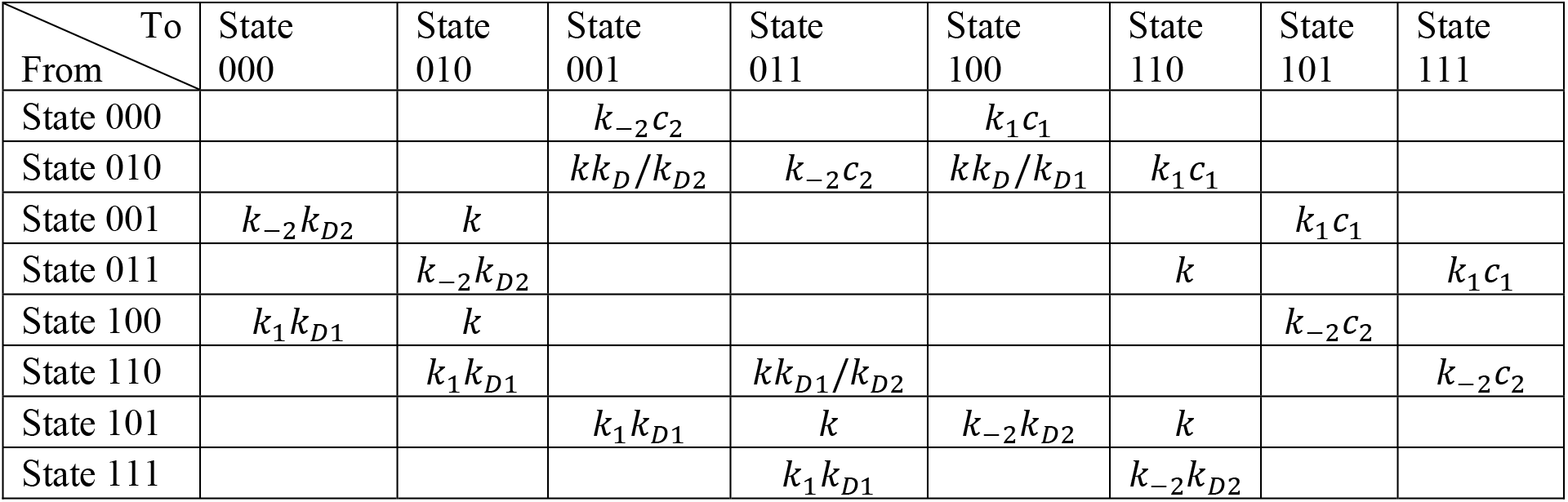
Transition rate constants

| From \ To | State 000 | State 010 | State 001 | State 011 | State 100 | State 110 | State 101 | State 111 |
| --- | --- | --- | --- | --- | --- | --- | --- | --- |
| State 000 | | | $k_{-2}c_2$ | | $k_1c_1$ | | | |
| State 010 | | | $kk_D/k_{D2}$ | $k_{-2}c_2$ | $kk_D/k_{D1}$ | $k_1c_1$ | | |
| State 001 | $k_{-2}k_{D2}$ | $k$ | | | | | $k_1c_1$ | |
| State 011 | | $k_{-2}k_{D2}$ | | | | $k$ | | $k_1c_1$ |
| State 100 | $k_1k_{D1}$ | $k$ | | | | | $k_{-2}c_2$ | |
| State 110 | | $k_1k_{D1}$ | | $kk_{D1}/k_{D2}$ | | | | $k_{-2}c_2$ |
| State 101 | | | $k_1k_{D1}$ | $k$ | $k_{-2}k_{D2}$ | $k$ | | |
| State 111 | | | | $k_1k_{D1}$ | | $k_{-2}k_{D2}$ | | |

### Experimental data vs. computational predictions

In order to validate our PMF-based theoretical-computational study, we performed 20 sets of *in vitro* experiments on human erythrocytes at four different temperatures between 5°C and 37°C for a wide range of glycerol concentrations (tabulated in Table IV). Five sets of data at 23°C are shown in Fig. 5 and all the other data sets are included in SI, Figs. S2 to S4. Below we discuss in detail the experimental data and the perfect agreement with the theoretical-computational predictions.

**Table IV.** Experimental data of *k*_−2_

| EC [glycerol] | 25 mM | 50 mM | 100 mM | 200 mM | 400 mM |
| --- | --- | --- | --- | --- | --- |
| 36.6 °C | $1.3 \times 10^3$ /s·M | $1.3 \times 10^3$ /s·M | $1.3 \times 10^3$ /s·M | $1.3 \times 10^3$ /s·M | $1.3 \times 10^3$ /s·M |
| 23.5 °C | $1.05 \times 10^3$ /s·M | $1.05 \times 10^3$ /s·M | $1.05 \times 10^3$ /s·M | $1.05 \times 10^3$ /s·M | $1.0 \times 10^3$ /s·M |
| 16.1 °C | $0.85 \times 10^3$ /s·M | $0.9 \times 10^3$ /s·M | $0.9 \times 10^3$ /s·M | $0.9 \times 10^3$ /s·M | $0.9 \times 10^3$ /s·M |
| 5.3 °C | $0.8 \times 10^3$ /s·M | $0.8 \times 10^3$ /s·M | $0.8 \times 10^3$ /s·M | $0.8 \times 10^3$ /s·M | $0.8 \times 10^3$ /s·M |

**Fig. 5.**
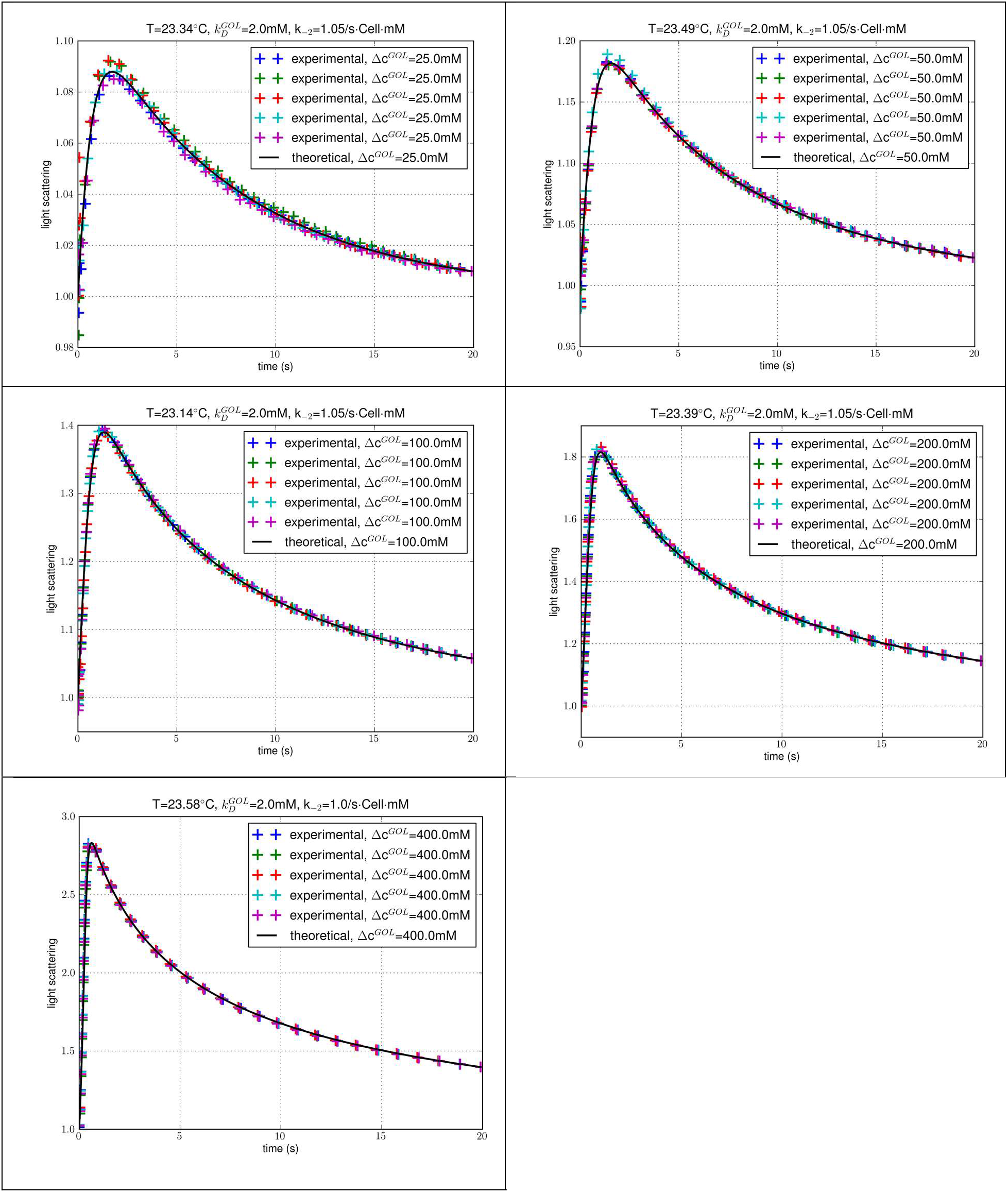
Experiments at ~23°C of glycerol uptake into human erythrocytes for extracellular glycerol concentration at 25 mM, 50 mM, 100 mM, 200 mM, and 400 mM. The data points (colored crosses) represent normalized intensity of light scattered at 90° immediately after mixing of an erythrocyte suspension in 0.7 ×PBS (containing no glycerol) with an equal volume of 0.7 ×PBS containing 2 × Δ*c*^*GOL*^ glycerol. Five colors represent five experimental repeats under identical conditions. The black solid curve is the predicated time course with a single fitting parameter *k*_−2_whose fitted values are shown. The fitting has a p-value less than 10^−5^ and a relative error ~5% in all five sets.

First of all, the fitted values of the single parameter *k*_−2_ turned out to be weakly dependent upon temperature and, for each temperature, nearly independent of the glycerol concentration ranging from 25 mM to 400 mM (Table IV). The theoretical-computational framework based on the PMF (Fig. 4) indeed captured all the significant factors of glycerol transport through AQP3. Otherwise, the fitted values of the fitting parameter would be strongly case-dependent.

For each given temperature, the time it took for the glycerol uptake to complete (i.e., for the IC glycerol concentration to reach the EC level) was nearly independent of the concentration gradient. It took approximately 20 seconds for the uptake of 25 mM glycerol to complete. Likewise, it took approximately 20 seconds for the uptake of 50 mM, 100 mM, 200 mM, or even 400 mM glycerol (Fig. 5 for experiments at ~23℃). The glycerol transport was clearly unsaturated. Otherwise, it would take a longer time for the uptake to complete at higher concentrations. The same conclusion can be drawn from experiments at 5℃ (~40 s for all of the five concentrations), 16℃ (~25 s for all of the five concentrations), 37℃ (~15 s for all of the five concentrations) (SI, Fig. S2 to S4). Indeed, glycerol transport through AQP3 is unsaturated in the concentration range up to ~1M in perfect agreement with the kinetics theory based on the PMF curve showing high AQP3-glycerol affinity at one of the three binding sites.

## CONCLUSIONS

From the large-scale simulation of human aquaglyceroporin AQP3 in erythrocyte membrane, we reached the time scale for direct observation of an event of transporting a glycerol molecule across the membrane and did observe such an event that two glycerol molecules moved collectively to transport one from the cytoplasm to the EC fluid. From the quantitative investigation, the kinetics theory based on the Gibbs free-energy gradient, the driving force for the transport process, we found the affinities of AQP3 for glycerol and the probabilities of glycerol transport pathways. With validation by experiments on human erythrocytes, we conclude that human aquaglyceroporin AQP3 has ~500/M affinity for glycerol and that AQP3 conducts glycerol transport in an unsaturable manner for glycerol concentrations <1M. Due to the glycerol-glycerol interaction facilitated by the AQP3 channel, an aquaglyceroporin having high affinity for its substrate can transport the substrate across the cell membrane in an unsaturated manner.

## Supplementary Information

Part I gives the long formulas of the transport kinetics. Part II gives four additional figures that are discussed but not included in the main text.

## Data availability

The Dataset (parameters, coordinates, scripts, etc.) to replicate this study is available at Harvard Dataverse, https://doi.org/10.7910/DVN/V9I4YQ.

## Author contributions

RAR and LYC conducted the theoretical-computational work; RC and HL did the experimental measurements; LYC conceptualized the research and wrote the paper; All participated in analyzing the data and editing the manuscript.

## Declaration

There are no conflicts to declare.

## Acknowledgements

The authors acknowledge use of computational time on Frontera at the Texas Advanced Computing Center at the University of Texas at Austin. Frontera is made possible by National Science Foundation award OAC-1818253.

## References

1. M. H. Jacobs, H. N. Glassman and K. Parpart Arthur, Journal of Cellular and Comparative Physiology, 1935, 7, 197–225.

2. P. G. LeFevre, The Journal of General Physiology, 1948, 31, 505.

3. W. D. Stein, Biochim. Biophys. Acta, 1962, 59, 47–65.

4. R. I. Macey, D. M. Karan and R. E. L. Farmer, in Biomembranes : Passive Permeability of Cell Membranes: A satellite symposium of the XXV Internationational Congress of Physiological Sciences, Munich, Germany, July 25–31, 1971, organized by the Department of Physiology, University of Nijmejen, Nijmejen, Netherlands, and held in Rotterdam, July 20–22, 1971, eds. F. Kreuzer and J. F. G. Slegers, Springer US, Boston, MA, 1972, DOI: 10.1007/978-1-4684-0961-1_22, pp. 331–340.

5. P. Mazur and R. H. Miller, Cryobiology, 1976, 13, 507–522.

6. A. Carlsen and J. O. Wieth, Acta Physiologica Scandinavica, 1976, 97, 501–513.

7. K. B. L. Heller, E. C. C.; Wilson, T. Hastings J. Bacteriol., 1980, 144, 5.

8. P. Agre, M. Bonhivers and M. J. Borgnia, J. Biol. Chem., 1998, 273, 14659–14662.

9. J. B. Heymann and A. Engel, Physiology, 1999, 14, 187–193.

10. K. Murata, K. Mitsuoka, T. Hirai, T. Walz, P. Agre, J. B. Heymann, A. Engel and Y. Fujiyoshi, Nature, 2000, 407, 599–605.

11. B. L. de Groot and H. Grubmüller, Science, 2001, 294, 2353–2357.

12. A. Engel and H. Stahlberg, in International Review of Cytology, ed. W. D. S. Thomas Zeuthen, Academic Press, Cambridge, MA, 2002, vol. 215, pp. 75–104.

13. T. Gonen and T. Walz, Quarterly Reviews of Biophysics, 2006, 39, 361–396.

14. J. M. Carbrey and P. Agre, ed. E. Beitz, Springer Berlin Heidelberg, 2009, vol. 190, pp. 3–28.

15. H. Sui, B.-G. Han, J. K. Lee, P. Walian and B. K. Jap, Nature, 2001, 414, 872–878.

16. G. Benga, European Biophysics Journal, 2013, 42, 33–46.

17. J. Abramson and A. S. Vartanian, Science, 2013, 340, 1294–1295.

18. A. S. Verkman, M. O. Anderson and M. C. Papadopoulos, Nat Rev Drug Discov, 2014, 13, 259–277.

19. A. Horner, F. Zocher, J. Preiner, N. Ollinger, C. Siligan, S. A. Akimov and P. Pohl, Science Advances, 2015, 1, e1400083.

20. D. Fu, A. Libson, L. J. W. Miercke, C. Weitzman, P. Nollert, J. Krucinski and R. M. Stroud, Science, 2000, 290, 481–486.

21. E. Tajkhorshid, P. Nollert, M. Ø. Jensen, L. J. W. Miercke, J. O’Connell, R. M. Stroud and K. Schulten, Science, 2002, 296, 525–530.

22. Z. E. Newby, J. O’Connell, 3rd, Y. Robles-Colmenares, S. Khademi, L. J. Miercke and R. M. Stroud, Nat Struct Mol Biol, 2008, 15, 619–625.

23. K. Ishibashi, S. Sasaki, K. Fushimi, S. Uchida, M. Kuwahara, H. Saito, T. Furukawa, K. Nakajima, Y. Yamaguchi and T. Gojobori, Proceedings of the National Academy of Sciences, 1994, 91, 6269.

24. M. Echevarría, E. E. Windhager and G. Frindt, J. Biol. Chem., 1996, 271, 25079–25082.

25. N. Roudier, J.-M. Verbavatz, C. Maurel, P. Ripoche and F. Tacnet, J. Biol. Chem., 1998, 273, 8407–8412.

26. T. Zeuthen and D. A. Klaerke, J. Biol. Chem., 1999, 274, 21631–21636.

27. B. Yang, T. Ma and A. S. Verkman, J. Biol. Chem., 2001, 276, 624–628.

28. N. Roudier, P. Ripoche, P. Gane, P. Y. Le Pennec, G. Daniels, J.-P. Cartron and P. Bailly, J. Biol. Chem., 2002, 277, 45854–45859.

29. M. Zelenina, S. Tritto, A. A. Bondar, S. Zelenin and A. Aperia, J. Biol. Chem., 2004, 279, 51939–51943.

30. H. Kida, T. Miyoshi, K. Manabe, N. Takahashi, T. Konno, S. Ueda, T. Chiba, T. Shimizu, Y. Okada and S. Morishima, The Journal of Membrane Biology, 2005, 208, 55–64.

31. K. Edashige, M. Tanaka, N. Ichimaru, S. Ota, K.-i. Yazawa, Y. Higashino, M. Sakamoto, Y. Yamaji, T. Kuwano, D. M. Valdez, F. W. Kleinhans and M. Kasai, Biology of Reproduction, 2006, 74, 625–632.

32. E. W. Miller, B. C. Dickinson and C. J. Chang, Proceedings of the National Academy of Sciences, 2010, 107, 15681–15686.

33. E. Campos, T. F. Moura, A. Oliva, P. Leandro and G. Soveral, Biochem. Biophys. Res. Commun., 2011, 408, 477–481.

34. G. Calamita, P. Gena, D. Ferri, A. Rosito, A. Rojek, S. Nielsen, R. A. Marinelli, G. Frühbeck and M. Svelto, Biology of the Cell, 2012, 104, 342–351.

35. A. P. Martins, A. Marrone, A. Ciancetta, A. Galán Cobo, M. Echevarría, T. F. Moura, N. Re, A. Casini and G. Soveral, PLoS ONE, 2012, 7, e37435.

36. D. C. Soler, X. Bai, L. Ortega, T. Pethukova, S. T. Nedorost, D. L. Popkin, K. D. Cooper and T. S. McCormick, Medical Hypotheses, 2015, 84, 498–503.

37. A. Spinello, A. de Almeida, A. Casini and G. Barone, J. Inorg. Biochem., 2016, 160, 78–84.

38. V. Graziani, A. Marrone, N. Re, C. Coletti, J. A. Platts and A. Casini, Chemistry – A European Journal, 2017, 23, 13802–13813.

39. M. Palmgren, M. Hernebring, S. Eriksson, K. Elbing, C. Geijer, S. Lasič, P. Dahl, J. S. Hansen, D. Topgaard and K. Lindkvist-Petersson, The Journal of Membrane Biology, 2017, 250, 629–639.

40. M. Arif, P. Kitchen, M. T. Conner, E. J. Hill, D. Nagel, R. M. Bill, S. J. Dunmore, A. L. Armesilla, S. Gross, A. R. Carmichael, A. C. Conner and J. E. Brown, Oncology Letters, 2018, 16, 713–720.

41. G. Calamita, J. Perret and C. Delporte, Frontiers in Physiology, 2018, 9, 851.

42. Z. Zhu, L. Jiao, T. Li, H. Wang, W. Wei and H. Qian, Oncology Letters, 2018, 16, 2661–2667.

43. A. P. Martins, A. Ciancetta, A. de Almeida, A. Marrone, N. Re, G. Soveral and A. Casini, ChemMedChem, 2013, 8, 1086–1092.

44. C. Maurel, J. Reizer, J. I. Schroeder, M. J. Chrispeels and M. H. Saier, J. Biol. Chem., 1994, 269, 11869–11872.

45. M. Hansen, J. F. J. Kun, J. E. Schultz and E. Beitz, J. Biol. Chem., 2002, 277, 4874–4882.

46. M. Ishii, K. Ohta, T. Katano, K. Urano, J. Watanabe, A. Miyamoto, K. Inoue and H. Yuasa, Cell. Physiol. Biochem., 2011, 27, 749–756.

47. Y. Ohgusu, K.-y. Ohta, M. Ishii, T. Katano, K. Urano, J. Watanabe, K. Inoue and H. Yuasa, Drug Metabolism and Pharmacokinetics, 2008, 23, 279–284.

48. A. Delgado-Bermúdez, M. Llavanera, L. Fernández-Bastit, S. Recuero, Y. Mateo-Otero, S. Bonet, I. Barranco, B. Fernández-Fuertes and M. Yeste, Journal of animal science and biotechnology, 2019, 10, 77–77.

49. A. Delgado-Bermúdez, M. Llavanera, S. Recuero, Y. Mateo-Otero, S. Bonet, I. Barranco, B. Fernandez-Fuertes and M. Yeste, International Journal of Molecular Sciences, 2019, 20, 6255.

50. A. Delgado-Bermúdez, F. Noto, S. Bonilla-Correal, E. Garcia-Bonavila, J. Catalán, M. Papas, S. Bonet, J. Miró and M. Yeste, Biology, 2019, 8, 85.

51. J. A. Freites, K. L. Németh-Cahalan, J. E. Hall and D. J. Tobias, Biochimica et Biophysica Acta (BBA) - Biomembranes, 2019, 1861, 988–996.

52. E. E. S. Ong and J.-L. Liow, Fluid Phase Equilib., 2019, 481, 55–65.

53. R. A. Rodriguez, H. Liang, L. Y. Chen, G. Plascencia-Villa and G. Perry, Biochimica et Biophysica Acta (BBA) - Biomembranes, 2019, 1861, 768–775.

54. J. Rodríguez-Gamir, J. Xue, M. J. Clearwater, D. F. Meason, P. W. Clinton and J.-C. Domec, Plant, Cell & Environment, 2019, 42, 717–729.

55. M. Sisto, D. Ribatti and S. Lisi, in Advances in Protein Chemistry and Structural Biology, ed. R. Donev, Academic Press, 2019, vol. 116, pp. 311–345.

56. Y. Sonntag, P. Gena, A. Maggio, T. Singh, I. Artner, M. K. Oklinski, U. Johanson, P. Kjellbom, J. D. Nieland, S. Nielsen, G. Calamita and M. Rützler, J. Biol. Chem., 2019.

57. O. H. Wittekindt and P. Dietl, Pflugers Archiv : European journal of physiology, 2019, 471, 519–532.

58. A. Zaragoza, M. A. Gonzalez, L. Joly, I. López-Montero, M. A. Canales, A. L. Benavides and C. Valeriani, PCCP, 2019, 21, 13653–13667.

59. X. Zhou and F. Zhu, Journal of Chemical Information and Modeling, 2019, 59, 777–785.

60. S. W. de Maré, R. Venskutonytė, S. Eltschkner, B. L. de Groot and K. Lindkvist-Petersson, Structure, 2020, 28, 215–222.e213.

61. M. Hernando, G. Orriss, J. Perodeau, S. Lei, F. G. Ferens, T. R. Patel, J. Stetefeld, A. J. Nieuwkoop and J. D. O’Neil, Biochimica et Biophysica Acta (BBA) - Biomembranes, 2020, 1862, 183191.

62. T. Katano, Y. Ito, K. Ohta, T. Yasujima, K. Inoue and H. Yuasa, Drug Metabolism and Pharmacokinetics, 2014, 29, 244–248.

63. Y. Liu, D. Promeneur, A. Rojek, N. Kumar, J. Frøkiær, S. Nielsen, L. S. King, P. Agre and J. M. Carbrey, Proceedings of the National Academy of Sciences, 2007, 104, 12560–12564.

64. H. Tsukaguchi, C. Shayakul, U. V. Berger, B. Mackenzie, S. Devidas, W. B. Guggino, A. N. van Hoek and M. A. Hediger, J. Biol. Chem., 1998, 273, 24737–24743.

65. S. Hatakeyama, Y. Yoshida, T. Tani, Y. Koyama, K. Nihei, K. Ohshiro, J.-I. Kamiie, E. Yaoita, T. Suda, K. Hatakeyama and T. Yamamoto, Biochem. Biophys. Res. Commun., 2001, 287, 814–819.

66. K. Ishibashi, T. Morinaga, M. Kuwahara, S. Sasaki and M. Imai, Biochimica et Biophysica Acta (BBA) - Gene Structure and Expression, 2002, 1576, 335–340.

67. A. Mobasheri, M. Shakibaei and D. Marples, Histochemistry and Cell Biology, 2004, 121, 463–471.

68. D. C. Soler, A. E. Young, A. D. Griffith, P. F. Fu, K. D. Cooper, T. S. McCormick and D. L. Popkin, Experimental dermatology, 2017, 26, 949–951.

69. K. Gotfryd, A. F. Mósca, J. W. Missel, S. F. Truelsen, K. Wang, M. Spulber, S. Krabbe, C. Hélix-Nielsen, U. Laforenza, G. Soveral, P. A. Pedersen and P. Gourdon, Nature Communications, 2018, 9, 4749.

70. E. Beitz, S. Pavlovic-Djuranovic, M. Yasui, P. Agre and J. E. Schultz, Proc Natl Acad Sci U S A, 2004, 101, 1153–1158.

71. K. Edashige, S. Ohta, M. Tanaka, T. Kuwano, D. M. Valdez, Jr., T. Hara, B. Jin, S.-i. Takahashi, S. Seki, C. Koshimoto and M. Kasai, Biology of Reproduction, 2007, 77, 365–375.

72. J. M. Lahmann, J. D. Benson and A. Z. Higgins, Cryobiology, 2018, 80, 1–11.

73. M. J. Borgnia and P. Agre, Proceedings of the National Academy of Sciences, 2001, 98, 2888–2893.

74. L. Y. Chen, Journal of Structural Biology, 2013, 181, 71–76.

75. L. Y. Chen, Biochimica Et Biophysica Acta-Biomembranes, 2013, 1828, 1786–1793.

76. A. Vahedi-Faridi, Lodowski, D., Kowatz, T., PDB, 2020, DOI: 10.2210/pdb6n1g/pdb.

77. A. H. Bryk and J. R. Wiśniewski, Journal of Proteome Research, 2017, 16, 2752–2761.

78. J. T. Dodge and G. B. Phillips, Journal of Lipid Research, 1967, 8, 667–675.

79. M. S. Bretscher, Nature: New biology, 1972, 236, 11–12.

80. R. F. A. Zwaal and A. J. Schroit, Blood, 1997, 89, 1121.

81. J. A. Virtanen, K. H. Cheng and P. Somerharju, Proceedings of the National Academy of Sciences, 1998, 95, 4964.

82. R. Chan, M. Falato, H. Liang and L. Y. Chen, RSC Advances, 2020, 10, 21283–21291.

83. W. Humphrey, A. Dalke and K. Schulten, Journal of Molecular Graphics, 1996, 14, 33–38.

84. S. Jo, T. Kim, V. G. Iyer and W. Im, J. Comput. Chem., 2008, 29, 1859–1865.

85. J. C. Phillips, R. Braun, W. Wang, J. Gumbart, E. Tajkhorshid, E. Villa, C. Chipot, R. D. Skeel, L. Kalé and K. Schulten, J. Comput. Chem., 2005, 26, 1781–1802.

86. R. B. Best, X. Zhu, J. Shim, P. E. M. Lopes, J. Mittal, M. Feig and A. D. MacKerell, Journal of Chemical Theory and Computation, 2012, 8, 3257–3273.

87. K. Vanommeslaeghe, E. Hatcher, C. Acharya, S. Kundu, S. Zhong, J. Shim, E. Darian, O. Guvench, P. Lopes, I. Vorobyov and A. D. Mackerell, J. Comput. Chem., 2010, 31, 671–690.

88. J. B. Klauda, R. M. Venable, J. A. Freites, J. W. O’Connor, D. J. Tobias, C. Mondragon-Ramirez, I. Vorobyov, A. D. MacKerell and R. W. Pastor, The Journal of Physical Chemistry B, 2010, 114, 7830–7843.

89. L. D. Landau and E. M. Lifshitz, Statistical Physics, Part 1, Pergamon Press, Tarrytown, 1985.

90. H.-J. Woo and B. Roux, Proc. Natl. Acad. Sci. U.S.A., 2005, 102, 6825–6830.

91. R. A. Rodriguez, L. Yu and L. Y. Chen, J Chem Theory Comput, 2015, 11 4427–4438.

