## Supplemental Information for "Quantitative study of unsaturated transport of glycerol through aquaglyceroporin that has high affinity for glycerol"

In Part I, first part of this supplemental information, we give the long formulas. In Part II, the second part of this SI, we provide four additional figures that are discussed but not included in the main text.

### Part I.

The probabilities for the seven states are governed by the following equations of transport kinetics:

$$\begin{aligned}
\frac{dp_{000}}{dt} &= k_1\, k_{D1}\, p_{100} + k_{-2}\, k_{D2}\, p_{001} - (k_1\, c_1 + k_{-2}\, c_2)\, p_{000},\\
\frac{dp_{100}}{dt} &= k_1\, c_1\, p_{000} - k_1\, k_{D1}\, p_{100} + k\frac{k_D}{k_{D1}} p_{010} - k\, p_{100},\\
\frac{dp_{010}}{dt} &= k\, p_{100} + k\, p_{001} + k_1\, k_{D1}\, p_{110} + k_{-2}\, k_{D2}\, p_{011} - \left(\frac{k\, k_D}{k_{D1}} + \frac{k\, k_D}{k_{D2}}\right) p_{010} - (k_1\, c_1 + k_{-2}\, c_2)\, p_{010},\\
\frac{dp_{001}}{dt} &= k_{-2} c_2 p_{000} - k_{-2}\, k_{D2}\, p_{001} + k\frac{k_D}{k_{D2}} p_{010} - k\, p_{001},\\
\frac{dp_{110}}{dt} &= k_1\, c_1\, p_{010} + k_{-2}\, k_{D2}\, p_{111} + k\, p_{011} - (k\,\frac{k_{D1}}{k_{D2}} + k_1\, k_{D1} + k_{-2}\, c_2)\, p_{110},\\
\frac{dp_{011}}{dt} &= k_{-2}\, c_2\, p_{010} + k_1\, k_{D1}\, p_{111} + k\,\frac{k_{D1}}{k_{D2}}\, p_{110} - (k + k_1\, c_1 + k_{-2}\, k_{D2})\, p_{011},\\
\frac{dp_{111}}{dt} &= k_{-2}\, c_2\, p_{110} + k_1\, c_1\, p_{011} - (k_1\, k_{D1} + k_{-2}\, k_{D2})\, p_{111}.
\end{aligned} \tag{1}$$

Here the rate constants are illustrated in Fig. 3 of the main text. It can be verified that these probabilities are normalizable:

$$p_{000} + p_{100} + p_{010} + p_{001} + p_{011} + p_{110} + p_{111} = 1. \tag{2}$$

The flux going into the IC side and the flux leaving the EC side are, respectively,

$$\begin{aligned}
J_2 &= k_{-2}\, k_{D2}\, p_{001} - k_{-2}\, c_2\, p_{000} + k_{-2}\, k_{D2}\, p_{011} - k_{-2}\, c_2\, p_{010} + k_{-2}\, k_{D2}\, p_{111} - k_{-2}\, c_2\, p_{110},\\
J_1 &= -k_1 k_{D1}\, p_{111} + k_1\, c_1\, p_{011} + k_1\, c_1\, p_{010} - k_1\, k_{D1}\, p_{110} + k_1\, c_1\, p_{000} - k_1\, k_{D1}\, p_{100}.
\end{aligned} \tag{3}$$

Under the quasi-stationary conditions, we found $J = J_2 = J_1 = K(c_e - c)$ with the EC concentration noted as $c_2 = c_e$ and the IC concentration $c_1 = c$. The rate coeffient $K$ is a very complex functions of the two concentrations as follows:

$$
\begin{aligned}
K = {} & \\
& (k\, k1\, k2r\, kD1\, kD2\, (c2^2\, k2r^2\, (k + k1\, kD1) + c1^2\, k1^2\, (k + k2r\, kD2) + kD\, (k1\, kD1 + k2r\, kD2)\, (k\, (k1 + k2r) + k1\, k2r\, kD2) + \\
& \quad c1\, k1\, (k\, (k1\, kD1 + k2r\, kD2) + c2\, k2r\, (2\, k + k1\, kD1 + k2r\, kD2) + k2r\, kD2\, (k1\, (kD + kD1) + k2r\, kD2)) + \\
& \quad c2\, k2r\, (k\, (k1\, kD1 + k2r\, kD2) + k1\, (k1\, kD1^2 + k2r\, (kD + kD1)\, kD2)))) \,/ \\
& (c2^3\, k2r^3\, (k + k1\, kD1)^2\, kD2 + c1^3\, k1^3\, (k + k2r\, kD2)\, (k\, kD1 + k2r\, kD2\, (c2 + kD2)) + \\
& \quad kD\, kD1\, kD2\, (k1\, kD1 + k2r\, kD2)\, (k\, (k1 + k2r) + k1\, k2r\, kD2)\, (k\, (k1 + k2r) + k1\, k2r\, (kD1 + kD2)) + \\
& \quad c2^2\, k2r^2\, (k + k1\, kD1)\, (k\, (kD1 + kD2)\, (k1\, kD1 + k2r\, kD2) + k1\, kD2\, (k1\, kD1^2 + 2\, k2r\, kD1\, kD2 + k2r\, kD\, (kD1 + kD2))) + \\
& \quad c1^2\, k1^2\, (c2^2\, k2r^2\, kD2\, (2\, k + k1\, kD1 + k2r\, kD2) + \\
& \qquad c2\, k2r\, (k^2\, (2\, kD1 + kD2) + k\, k1\, kD1\, (kD1 + 2\, kD2) + k\, k2r\, kD2\, (kD1 + 4\, kD2) + k2r\, kD2^2\, (3\, k1\, kD1 + 2\, k2r\, kD2)) + \\
& \qquad (k + k2r\, kD2)\, (k\, (kD1 + kD2)\, (k1\, kD1 + k2r\, kD2) + k2r\, kD2\, (2\, k1\, kD1\, kD2 + k2r\, kD2^2 + k1\, kD\, (kD1 + kD2)))) + \\
& \quad c2\, k2r\, (k^2\, (k1 + k2r)\, (k1\, kD1 + k2r\, kD2)\, (kD1\, kD2 + kD\, (kD1 + kD2)) + \\
& \qquad k1^2\, k2r\, kD1\, kD2\, (k1\, kD1\, (kD1\, kD2 + kD\, (kD1 + kD2)) + k2r\, kD2\, (kD1\, kD2 + 2\, kD\, (kD1 + kD2))) + \\
& \qquad k\, k1\, (k1^2\, kD1^2\, (kD1\, kD2 + kD\, (kD1 + kD2)) + k2r^2\, kD2\, (kD1\, kD2\, (kD1 + kD2) + kD\, (kD1^2 + 3\, kD1\, kD2 + kD2^2)) + \\
& \qquad\quad k1\, k2r\, kD1\, (kD1\, kD2\, (kD1 + 2\, kD2) + kD\, (kD1^2 + 3\, kD1\, kD2 + 3\, kD2^2)))) + \\
& \quad c1\, k1\, (c2^3\, k2r^3\, (k + k1\, kD1)\, kD2 + k^2\, (k1 + k2r)\, (k1\, kD1 + k2r\, kD2)\, (kD1\, kD2 + kD\, (kD1 + kD2)) + \\
& \qquad c2^2\, k2r^2\, (k + k1\, kD1)\, (k\, (kD1 + 2\, kD2) + kD2\, (2\, k1\, kD1 + 3\, k2r\, kD2)) + \\
& \qquad k1\, k2r^2\, kD2^2\, (k2r\, kD2\, (kD1\, kD2 + kD\, (kD1 + kD2)) + k1\, kD1\, (kD1\, kD2 + 2\, kD\, (kD1 + kD2))) + \\
& \qquad k\, k2r\, kD2\, (k1^2\, kD1\, (2\, kD\, kD1 + 3\, kD\, kD2 + 2\, kD1\, kD2) + k2r^2\, kD2\, (kD1\, kD2 + kD\, (kD1 + kD2)) + \\
& \qquad\quad k1\, k2r\, (kD1\, kD2\, (kD1 + 2\, kD2) + kD\, (kD1^2 + 4\, kD1\, kD2 + 2\, kD2^2))) + \\
& \qquad c2\, k2r\, (2\, k^2\, (kD1 + kD2)\, (k1\, kD1 + k2r\, kD2) + k1\, k2r\, kD2\, (k1\, kD1 + k2r\, kD2)\, (3\, kD1\, kD2 + kD\, (kD1 + kD2)) + \\
& \qquad\quad k\, (k1^2\, kD1^2\, (kD1 + 2\, kD2) + k2r^2\, kD2^2\, (kD1 + 2\, kD2) + 2\, k1\, k2r\, kD2\, (kD\, (kD1 + kD2) + kD1\, (kD1 + 3\, kD2))))))
\end{aligned}
\tag{4}
$$

where $c1 = c_e$, $c2 = c$ and $k2r = k_{-2}$ were adopted in the main text.

**Part II.**

Fig. S1 illustrates the all-atom model of an AQP3 tetramer embedded in a patch of lipid bilayers mimicking the lipid compositions of the inner and the outer leaflets of human erythrocyte membrane.

Fig. S2 to S4 show the experimental data along with the theoretical curves for glycerol uptake at ~5°C, ~16°C, and ~37°C respectively.

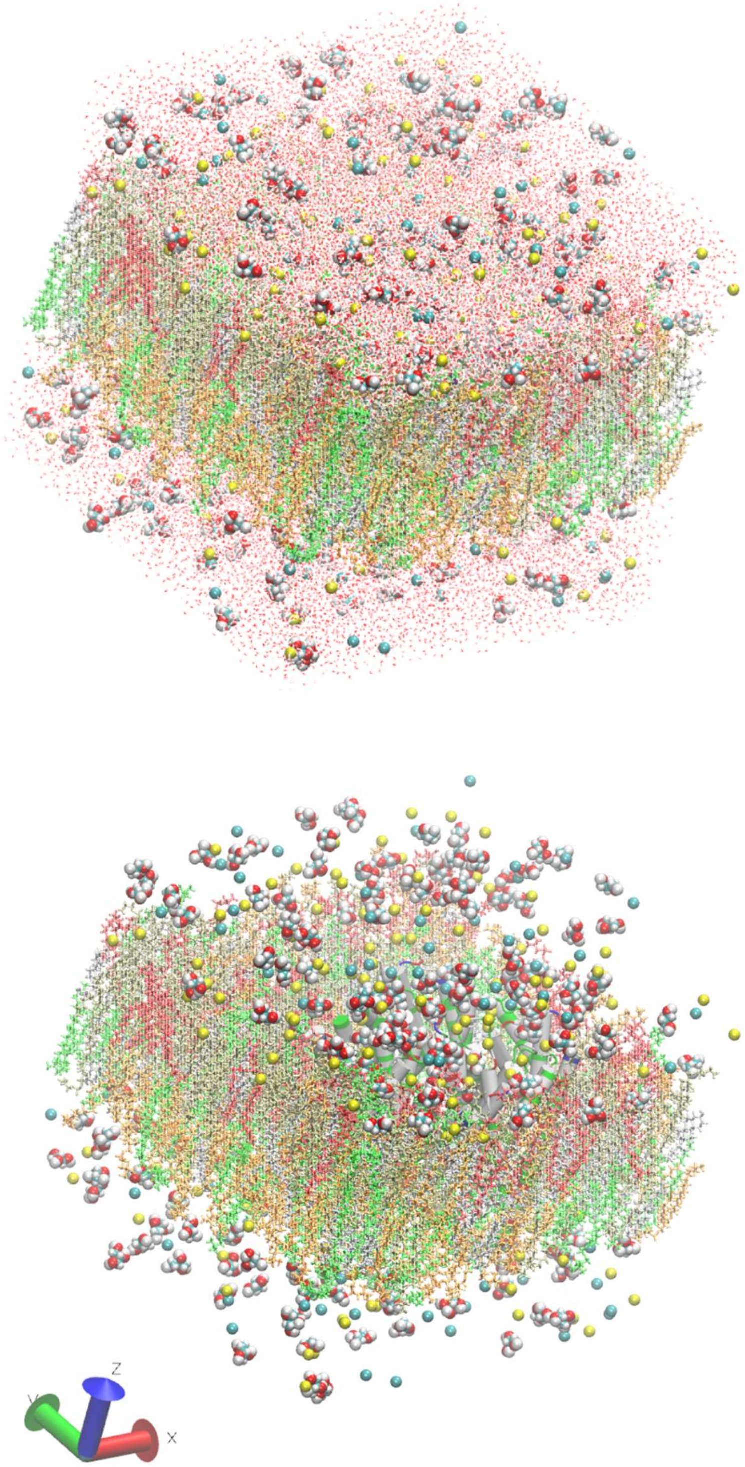


**Fig. S1.** All-atom model system of an AQP3 tetramer in a membrane patch mimicking the lipid composition of the human erythrocyte membrane. The system consists of 156,137 atoms. The intracellular space is located at $z > 20$Å and the extracellular space at $z < -20$Å. The water molecules are shown in the top panel as red-and-white dots but not shown in the bottom panel for clearer views of all the other constituents of the system. The glycerol molecules and salt (NaCl) are shown as spheres colored by atom names (C, cyan; O, red; N, blue; H, white; Cl, cyan; K, metallic; Na, yellow). The lipids are represented as licorices colored lipid names (POPC, orange; POPE, tan; POPS, red; SSM, green; CHL, silver). The proteins are presented as surfaces colored by residue types (hydrophilic, green; hydrophobic, white; positively charged, blue; negatively charged, red). Molecular graphics in this paper were rendered with VMD[1].

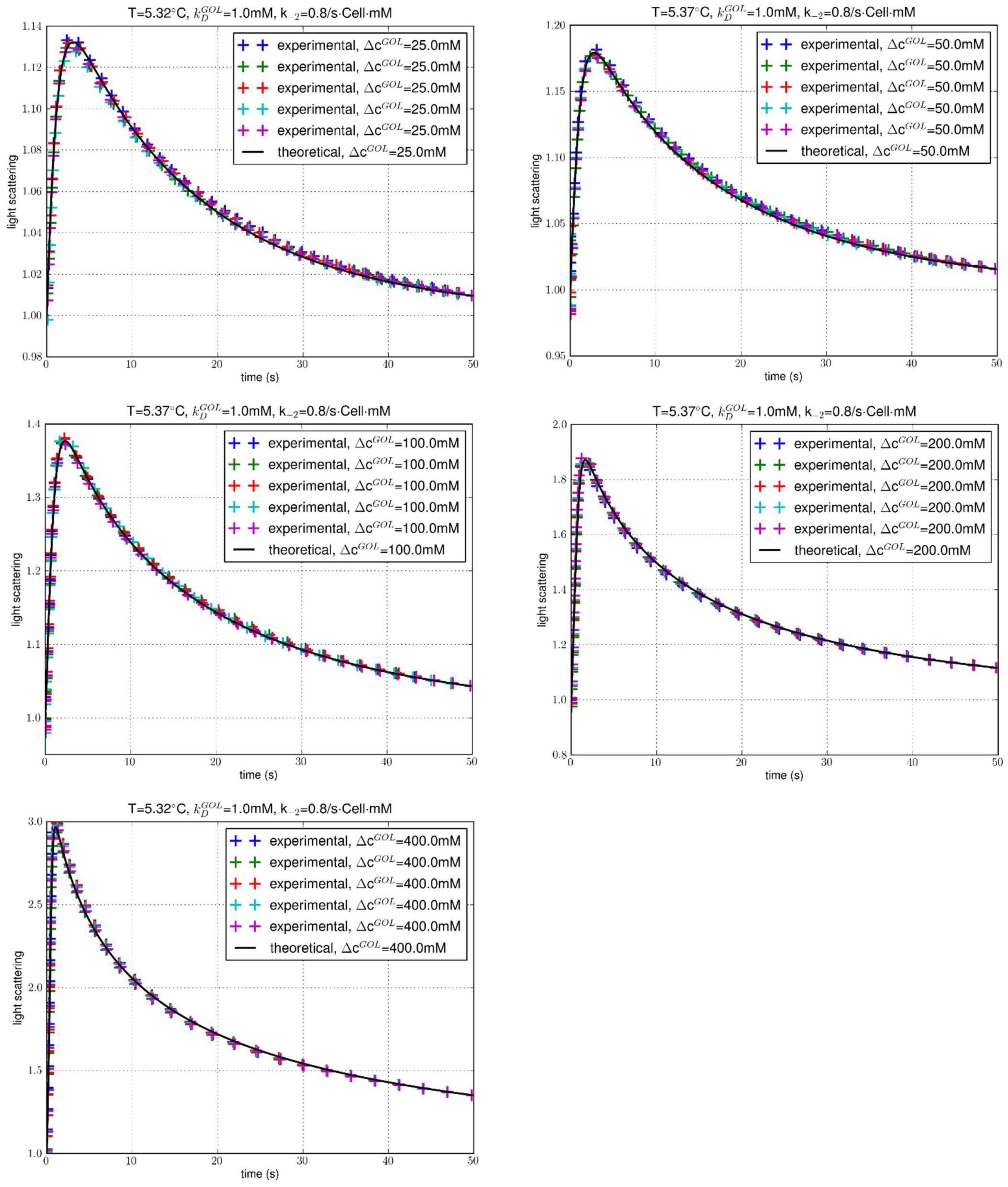


**Fig. S2.** Experiments at ~5°C of glycerol uptake into human erythrocytes for extracellular glycerol concentration at 25 mM, 50 mM, 100 mM, 200 mM, and 400 mM. The data points (colored crosses) represent normalized intensity of light scattered at 90° immediately after mixing of an erythrocyte suspension in $0.7$ ×PBS (containing no glycerol) with an equal volume of $0.7$ ×PBS containing $2 \times \Delta c^{GOL}$ glycerol. Five colors represent five experimental repeats under identical conditions. The black solid curve is the predicated time course with a single fitting parameter $k_{-2}$whose fitted values are shown. The fitting has a p-value less than $10^{-5}$ in all five sets and a relative error ~5%.

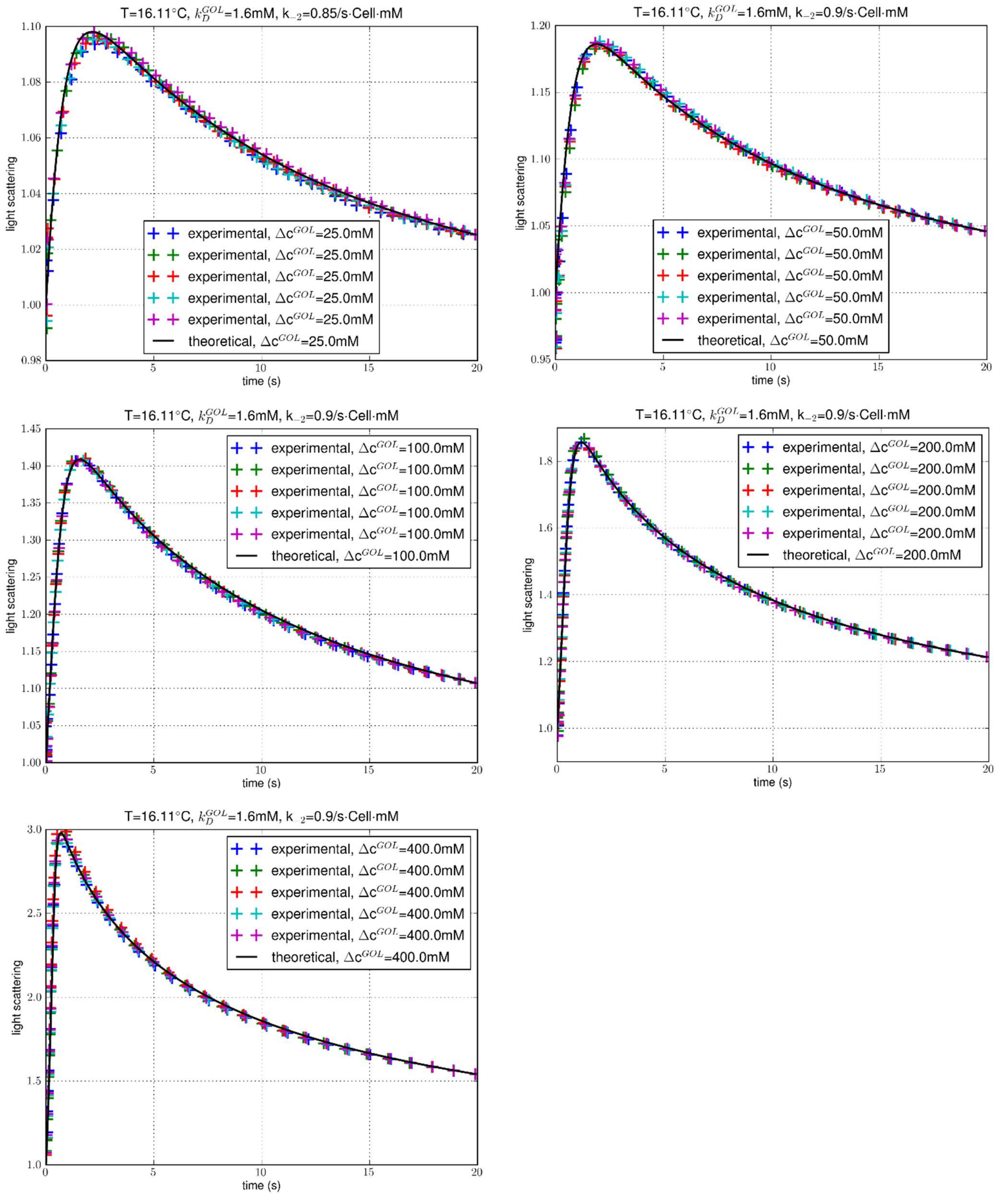


**Fig. S3.** Experiments at ~16°C of glycerol uptake into human erythrocytes for extracellular glycerol concentration at 25 mM, 50 mM, 100 mM, 200 mM, and 400 mM. The data points (colored crosses) represent normalized intensity of light scattered at 90° immediately after mixing of an erythrocyte suspension in $0.7$ ×PBS (containing no glycerol) with an equal volume of $0.7$ ×PBS containing $2 \times \Delta c^{GOL}$ glycerol. Five colors represent five experimental repeats under identical conditions. The black solid curve is the predicated time course with a single fitting parameter $k_{-2}$whose fitted values are shown. The fitting has a p-value less than $10^{-5}$ in all five sets and a relative error ~5%.

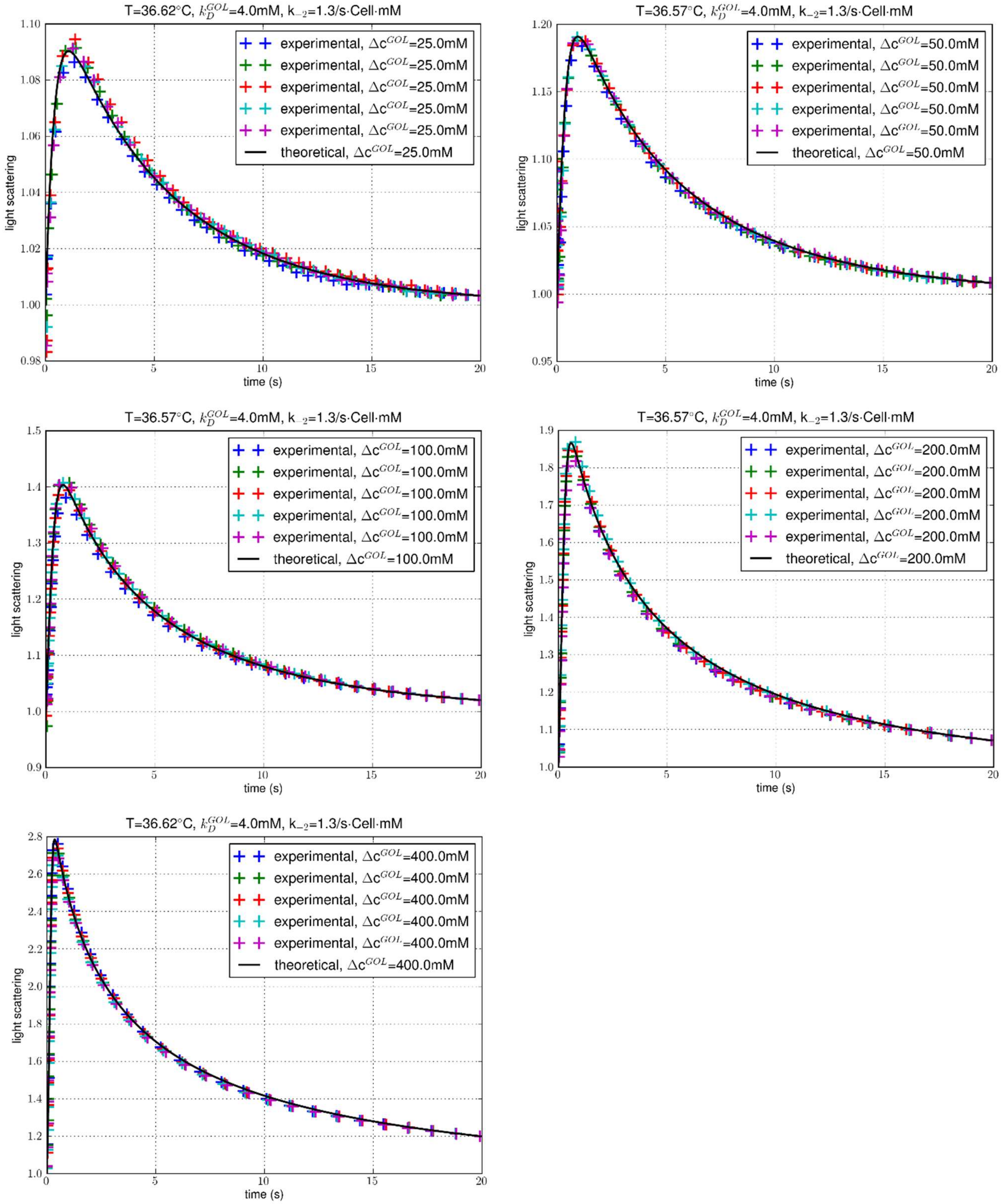


**Fig. S4**. Experiments at ~37°C of glycerol uptake into human erythrocytes for extracellular glycerol concentration at 25 mM, 50 mM, 100 mM, 200 mM, and 400 mM. The data points (colored crosses) represent normalized intensity of light scattered at 90° immediately after mixing of an erythrocyte suspension in $0.7\ \times$PBS (containing no glycerol) with an equal volume of $0.7\ \times$PBS containing $2 \times \Delta c^{GOL}$ glycerol. Five colors represent five experimental repeats under identical conditions. The black solid curve is the predicated time course with a single fitting parameter $k_{-2}$whose fitted values are shown. The fitting has a p-value less than $10^{-5}$ in all five sets and a relative error ~5%.
